# Pyrimethamine and a potent analogue WCDD115 inhibit NRF2 by suppressing DHFR and one-carbon metabolism

**DOI:** 10.1101/2025.02.13.637433

**Authors:** Julius Chembo, Brittany M. Bowman, Kyle Lapak, Emily Wilkerson, Chorlada Paiboonrungruang, Kevin Cho, Matthew R. Medcalf, Gary J. Patti, Roland E. Dolle, Xiaoxin Chen, Paul Zolkind, Michael B. Major

**Author notes:** These authors contributed equally to this work (Julius Chembo, Brittany M. Bowman). Corresponding Author: Michael B. Major, Address: Washington University in St. Louis, 660 Euclid Avenue, Box 8228, St. Louis, MO 63110.

## Abstract

Nuclear factor erythroid 2-related factor 2 (NFE2L2/NRF2) is a critical mediator of the cellular oxidative stress response. Aberrant activation of NRF2 is common in lung and upper aerodigestive cancers, where it promotes tumor initiation and progression and confers resistance to chemotherapy, radiation therapy, and immune checkpoint inhibitors. As such, NRF2 therapeutic inhibitors are actively being sought. We previously reported that the antiparasitic drug Pyrimethamine (PYR) inhibits NRF2 in cell lines and in a NRF2-inducible genetically engineered mouse model. Here we design, synthesize, and define structure-activity relationships across a series of 25 PYR-based derivatives to reveal WCDD115 as a 22-fold more potent inhibitor of NRF2 (57nM versus 1.2µM). PYR is known to inhibit plasmodial and human dihydrofolate reductase (DHFR). We found that WCDD115 inhibits hDHFR with 31-fold greater potency than PYR (144nM versus 4.49µM). Metabolomics showed strong similarities between PYR, WCDD115 and methotrexate. Genetic, pharmacological and metabolic epistasis studies reveal that DHFR inactivation is required for NRF2 suppression by WCDD115 and PYR. Global and targeted proteomics revealed overlapping profiles for WCDD115, PYR and methotrexate, including suppression of NRF2 oxidative stress response and activation of TP53 and the DNA damage response. Therefore, PYR and a novel potent derivative WCDD115 are effective, indirect inhibitors of NRF2 and its antioxidant functions. These data underscore the importance of one-carbon metabolism for the NRF2 signaling pathway and support a new therapeutic strategy to suppress NRF2-driven cancer biology.

## Introduction

Nuclear factor erythroid 2-related factor 2 (*NFE2L2,* hereafter referred to as NRF2) is a transcription factor that plays crucial roles in the cellular defense against oxidative and electrophilic stress [1–3]. In healthy cells and in the absence of stress, NRF2 protein levels are kept low through ubiquitin-dependent proteasomal degradation. Although several E3 ubiquitin ligases are known to govern NRF2 protein stability, the Kelch-like ECH-associated protein 1 (KEAP1) and Cullin3 (CUL3) ubiquitin ligase complex figures most prominently in NRF2 suppression and stress response [4, 5]. Specifically, reactive cysteine residues within KEAP1 sense oxidative/electrophilic stress, resulting in a poorly understood conformational change within the NRF2/KEAP1/CUL3 complex that ultimately inactivates the complex [6]. As such, newly transcribed and translated NRF2 then bypasses the inactive KEAP1/CUL3 complex to enter the nucleus, where it heterodimerizes with small musculoaponeurotic fibroblast (sMAF) proteins to activate the transcription of a battery of antioxidant, metabolic, and drug detoxification genes to restore cell health.

Evolution favored NRF2 as a means to abate oxidative and electrophilic stress, thus ensuring cellular health and resiliency in the face of oxidative phosphorylation, electrophilic metabolites and xenobiotics [7]. The evolutionary pressures of oncogenesis similarly create positive selection forces for NRF2-driven biologist. Broadly speaking, constitutive activation of NRF2 provides resilience to the transformed cell that would otherwise succumb to oxidative, metabolic, or immune-dependent cell death [8–10]. In human cancers of the lung, esophagus, head and neck, liver, and bladder, high levels of NRF2 activating mutations, NRF2 copy number amplifications or KEAP1/CUL3 loss-of-function mutations are observed. Independently of KEAP1/NRF2/CUL3 mutations, altered protein-protein interactions and KEAP1-inactivating posttranslational modifications also activate NRF2 in cancer, most notably in tissues of the liver, breast and kidney [9]. Though commonly activated in human cancer, genetically engineered mouse models show that NRF2 activation by itself results in pre-cancerous histologies, but not carcinoma [11–14]. When combined with other mutations such as TP53, CDKN2A, and KRAS, NRF2 activation promotes cancer initiation, early stages of tumor progression, and resistance to chemotherapy, radiation therapy, and immune checkpoint inhibitors[15].

Given the importance of NRF2 in the progression and therapeutic resistance, drug development pipelines have been active in the search for NRF2 inhibitors and synthetic-lethal strategies for NRF2-active cancer [16]. Brusatol and halofuginone are two clinically studied indirect NRF2 inhibitors that sensitize NRF2 active cancer cells to cytotoxic chemo and radiation therapy [17, 18]. These drugs suppress global protein translation, resulting in selective loss of short-lived proteins like NRF2. Our previous cell-based small molecule screens of NRF2-dependent transcription identified 12 lead compounds, including mitoxantrone and pyrimethamine (PYR) [19]. PYR, an FDA-approved medicine commonly used in the treatment of toxoplasmosis and chloroquine-resistant malaria, inhibits plasmodial dihydrofolate reductase (pDHFR). We and others found that like pDHFR, PYR also inhibits human and murine DHFR [20–22]. Using a *Sox2- Cre*; *LSL-Nrf2^E79Q/+^*GEMM, we demonstrated that PYR decreases NRF2^E79Q^-driven hyperplasia of esophageal epithelium, reduces expression of NRF2 and its target genes (eg. *GCLM*, *GCLC*), and inhibits cell proliferation (BrdU) with no observed weight loss, or liver, kidney or blood toxicity [19]. In contrast to brusatol and halofuginone, we found that PYR does not impact global proteostasis or translation.

PYR has received recent attention as a repurposed drug for cancer treatment, and with important differences from methotrexate [21–23]. A systematic literature review identified 14 articles reporting the dose and impact of PYR in mouse models of 10 cancer types [24]. Extending from these preclinical studies, PYR has been or is currently being tested in two phase 1 cancer clinical trials. First, using high-throughput small molecule screens, David Frank’s laboratory discovered PYR as an indirect inhibitor of STAT3 signaling [20, 23]. Inhibition of DHFR and folate metabolism were found to be critical for PYR suppression of STAT3. A completed clinical trial of PYR as a STAT3 inhibitor in chronic lymphocytic leukemia reported *in vivo* efficacy of STAT3 suppression, overall safety, and stable disease for 50% of the patients (NCT01066663) [23]. Our recent discovery that PYR inhibits NRF2 in cells and mice led to an ongoing phase 1 trial in HPV-negative head and neck squamous cell carcinoma (NCT05678438). Mechanistically, the anti-cancer effects of PYR, including STAT3 and NRF2 suppression, appear dependent on DHFR suppression, though the specifics of the underlying mechanism(s) remain unclear. In this study, we utilized structure-activity relationship (SAR) studies to investigate the connection between PYR, DHFR, and NRF2. We report compound WCDD115 as a 31-fold and 22-fold more potent derivative of PYR for DHFR and NRF2 suppression, respectively. Our data establishes a previously unappreciated role for one-carbon metabolism in regulating NRF2 activity. More broadly, WCDD115, PYR, and methotrexate global proteomic profiles position NRF2 suppression as one of several affected cellular pathways that may contribute therapeutic value to oncology.

## Materials and methods

### Cell culture

HEK293T and cancer cell lines NCI-H460, A549, H1299, H2170, H1792, H2122, OE21, and PC9 were purchased from ATCC. KYSE70, KYSE110, KYSE180, KYSE450 are from DSMZ (Germany), KYSE450-KEAP1KO and KYSE70-KEAP1KO cells were generated with CRISPR-Cas9. The mutational status of KEAP1 and NRF2 for each cell line is shown in Table S1. The reporter cell line H1299 NQO1-eYFP, with eYFP inserted into the first intron of the NQO1 gene, was kindly provided by Dr. Alon and the Kahn Protein Dynamics Group [25]. All cell lines were authenticated by short tandem repeat analysis (LabCorp, Genetica Cell Line Testing). Mycoplasma testing was performed regularly via the mycoplasma detection kit (Thermo Fischer Scientific, NC0447826). All cell lines were cultured in RPMI 1640 medium (Corning/Fisher, MT10040C) supplemented with 10% Fetal Bovine Serum (FBS), except for HEK29Ts, which were cultured in DMEM (Thermo Fischer Scientific, MT10013CV) supplemented with 10% FBS and maintained at 37C in a humidified atmosphere containing 5% CO2.

### Compounds and reagents

Pyrimethamine (Cayman/Neta Scientific: CAYM-16472-500), WCDD115 (See supplemental methods), Methotrexate (Sigma, A6770-25MG), Cycloguanil (Cayman/Neta, CAYM-16861-10), Pemetrexed (Sigma, SML1490-10MG), 50X Hypoxanthine and Thymidine (Sigma, H0137-10VL), Doxycycline (Thermo Fischer Scientific, BP26535), Menadione (Sigma, M5625-25G), Hypoxanthine (Thermo Fischer Scientific, AC122010050) and Thymidine (Thermo Fischer Scientific, AC226740050) and Dimethyl sulfoxide (Fisher, BP231-100). Chemical synthesis for WCDD115 and all PYR analogues are provided in supplemental methods.

### Western blot analysis

Following treatment with drugs, cell media was aspirated, cells were washed with Phosphate Buffered Saline (PBS) (Corning, MT21031CV) and harvested by scraping. RIPA lysis buffer (50 mM Tris-HCl, pH 7.4 (Invitrogen: 15568-025), 150 mM NaCl (Promega, V4221), 10% glycerol (Thermo Fischer Scientific, BPG33-1), 2 mM Ethylenediaminetetraacetic acid (EDTA) (Sigma, 03690- 100ML), 0.2% sodium deoxycholate (bioWORLD, SKU-40420018-2), 0.1% Sodium dodecyl sulfate (SDS) (Sigma, CAS-151-21-3), 1% NP-40-Alternative containing protease and phosphatase inhibitors (Thermo Fisher: PI78429 and PI78426) and Benzonase (Santa Cruz Biotechnology, sc- 202391) was added to extract the proteins. A bicinchoninic acid (BCA) (Thermo Fisher, PI23209) assay was used to quantify and normalize protein concentration. Dithiothreitol (DTT) (Sigma, D0632- 5G) was used as the reducing agent and PageRuler^TM^ plus ladder (Thermo Fischer, 226619). Equal amounts of protein were loaded. Protein separation by size was conducted by loading equal amounts of protein on a 4-12% polyacrylamide gel electrophoresis and running in MOPS running buffer (Boston Bioproducts, BP178). The separated proteins were then transferred to a nitrocellulose membrane (Thermo Fischer Scientific, PI88018) and then blocked with 5% milk (Carnation) for 1h. Afterwards, membranes were incubated at 4°C overnight (on a rotator) with primary antibodies diluted in 5% Bovine serum albumin (BSA) (Thermo Fischer Scientific, BP1600-100): NRF2 (Cell Signaling E5F1A, 1:1000), KEAP1 (Cell Signaling D6B12, 1:1,000), Vinculin/VCL (Santa Cruz sc25336, 1:3,000), DHFR (Cell Signaling 45710S, 1:1,000), GCLC (Abcam ab190685, 1:1,000), SLC7A11 (Cell Signaling D2M7A, 1:1000), NQO1 (Novus NB200-209, 1:1,000), HMOX (Abcam Ab13248, 1:1000) and Licor Revert^TM^ 700 Total Protein Stain Kits for Western Blot (LI-COR NC1572646/926-11016). Membranes were additionally incubated with antibodies conjugated to fluorescent dyes: solution (LI-COR IRDye 680, 800; 1:10,000 in 5% milk) for 1h at room temperature. Ultimately, changes in protein expression were visualized on the LICOR Fluorescence imager.

### Quantification of DHFR activity and reactive oxygen species

Dihydrofolate Reductase Assay Kit (Sigma, CS0340) was used to assess changes in recombinant DHFR activity following treatment with pyrimethamine, methotrexate, and WCDD115. The activity of DHFR was tested according to the product manual with the following considerations: The concentrations of DHFR used was 3x10-3 units. NADPH (60μM), and DHF (50μM) were kept constant while the drug concentrations were titrated to establish IC50. ROS levels were analyzed by utilizing an oxidation-sensitive dye (CellROX Green Reagent: Invitrogen, C10444) as recommended by the manufacturer. Cells undergoing treatment with drug were additionally treated with 20μM Menadione for 6 h to induce the ROS production. Cells were thereafter incubated with 5μM CellROX at 37°C for 1h. After aspiration of CellRox, phosphate-buffered saline was used to wash the cells. Imaging was done using the Incucyte S3 Live Cell Analysis System. ROS was quantified as a ratio of total green intensity to confluence mask.

### Cell line engineering

To generate recombinant lentivirus, a mixture containing poly-ethylenimine combined with the gene of interest, packaging psPAX2 (Addgene, catalog no. 12260) and envelope VSV-G plasmids (Addgene, catalog no. 12259) was made, and incubated at room temperature for 10 minutes. The mixture was then pipetted onto HEK293T cells. The virus was collected and filtered after 48 h. Cells were transduced with lentivirus for 24h with polybrene (8μg/ml) before the addition of antibiotics for positive selection. To generate DHFR knockout KYSE70 cells, we used sgRNAs targeting DHFR (CCDS47240) exon 3 (GAG-AAG-AAT-CGA-CCT-TTA-AA), and exon 4 (GAC-ATG-GTC-TGG-ATA-GTT-GG). These guides were cloned into the AarI restriction site of the VDB783 vector (kind gift from the McManus Lab) containing a puromycin resistance cassette. KYSE70 cells were first transduced with pLenti-Cas9 containing a blasticidin resistance cassette and then with the DHFR sgRNA vectors. After transduction, cells were selected with 2.5μg/mL blasticidin and 1μg/mL puromycin in the presence of hypoxanthine(H) and thymidine(T). Monoclonal populations were generated by single-cell dilutions in 96-well plates, again in the presence of HT. Genomic KO was confirmed by Sanger sequencing and western blot analysis. To generate doxycycline-inducible NRF2 shRNA KYSE70 cells, we obtained a dox-inducible short hairpin RNAs (shRNAs) targeting human NRF2 from Addgene: #136584, with a sequence of AGA-GCA-AGA-TTT-AGA-TCA-TTT- CTG-CAG-AAA-TGA-TCT-AAA-TCT-TGC-TCT. As a control, a shRNA targeting firefly luciferase was used (Addgene: #136587). KYSE70 cells were transduced with the virus for 24h in the presence of polybrene (8μg/mL). Post-transduction, the cells were cultured in media containing 1μg/mL puromycin.

### RNA Extraction and qRT-PCR

RNA extraction was performed using the TRIzol™ Reagent protocol (Thermo Fischer, 15596026), with RNAse-free water used for elution. RNA yield was determined using a NanoDrop One spectrophotometer (Thermo Fischer Scientific). Reverse transcription was carried out using the Invitrogen™ SuperScript™ IV VILO™ master mix kit (cat. no. 11766050). Quantitative real-time PCR was conducted with PowerUp™ SYBR™ Green Master Mix (Thermo Fischer, A25778) on a QuantStudio 6 Flex Real-Time PCR System (Thermo Fisher). mRNA levels were normalized to RPL13, and the comparative Ct method was employed to calculate fold changes in expression. Statistical significance was determined using a one-way ANOVA test.

### Sample preparation for mass spectrometry

Protein extraction, digestion and purifications were performed essentially as previously reported [26]. Briefly, cellular lysis was carried out using a urea-based lysis buffer containing 8 M urea, 70 mM NaCl, 1 mM EDTA, and 50 mM tris (pH 8.0) with addition of phosphatase and protease inhibitor cocktails (Halt, cat. no. 78429 and 78420). To normalize protein loading the BCA Protein Assay Kit (Thermo Fischer, 23225) was used. Subsequently, samples were treated with a reducing agent (5 mM DTT) at 37°C for 45 minutes and alkylated with chloroacetamide (50 mM) for 20 minutes at 25°C. The lysates underwent sequential digestion: first with LysC (20 mAu) at 30°C for 2 h, followed by trypsin (20μg/1mg protein) digestion at 37°C for 18 h. Before each digestion step, lysates were diluted to specific urea concentrations (3M before LysC, 1.5M before trypsin digest with 50 mM tris, pH 8.0). Trypsin activity was neutralized (1% formic acid), and the resulting peptides were desalted (Strata-X columns, 10 mg/ml; Phenomenex), dried using a speed vacuum, and resuspended for further analysis in 2% acetonitrile and 0.1% formic acid.

### Targeted protein mass spectrometry

For the targeted protein analyses of KYSE70 cells, a custom internal standard triggered-parallel reaction monitoring (OIS-PRM) method was used as previously reported [26]. OIS-PRM analyses report the ratios between pairs of unlabeled endogenous and Stable Isotope Labeled (SIL) internal standard peptides. The SIL peptides are catalogued in Table S1 of [26] and were injected at a nominal abundance of 40 fmol for every 1000 ng of endogenous peptide. Peak area ratios (PARs) were calculated as described previously but without the normalization procedure [26, 27].

### Data-dependent acquisition mass spectrometry

Tryptic peptides were separated by reversed-phase nanoflow chromatography using an Ultimate 3000 RSLCnano System (Thermo Fisher Scientific) with a uPAC Trapping column (Thermo Fischer, COL-TRPNANO16G1B2) and a 200 cm µPAC HPLC column (Thermo Fischer, COL- NANO200G1B). For peptide separation and elution, mobile phase A (MPA) was 0.1% formic acid (FA) in water and mobile phase B (MPB) was 0.1% FA in acetonitrile. Peptides were injected onto the trap column at 10 µL/min for 3 minutes using the loading pump. Initially the nanoflow rate was set at 0.7 µL/min and 2% mobile phase B while the peptides were loaded onto the trap column. A multi-step gradient was applied with a ramp from 2-8% MPB from 2.250-5 minutes, and 8-14.5% from 5-23 minutes. The flow rate was then dropped to 0.3 µL/min and the gradient was 14.5%-25% from 23-81.5 minutes, then 25-35% MPB from 81.5-101.5 minutes. The column was then washed using seesaw gradients up to 90% MPB followed by re-equilibration at 0.7 µL/min.

Mass spectrometry analysis was performed on a Thermo Fischer Orbitrap Eclipse. Full MS scans (375-2000 m/z) were acquired in the Orbitrap at 240K resolution with a 250% normalized automatic gain control (AGC), and auto max injection time. Precursors were selected for fragmentation using data-dependent top speed mode with a cycle time of 3 seconds and filtering for charge states 2-7. Precursors were isolated in the quadrupole with an isolation window of 0.7 m/z, fragmented by higher energy collisional dissociation (HCD) with a normalized collision energy of 35%, and analyzed in the ion trap at Turbo scan rate. After a precursor was selected for fragmentation and MS/MS analysis, the precursor with a 10ppm tolerance was excluded from repetitive sampling for 60 seconds. MS2 scans were collected with 100% normalized AGC and a maximum injection time of 35ms.

Raw data were processed using MaxQuant (version 2.1.1.0) [28]. Searches were performed against a target-decoy database of reviewed proteins (Uniprot (human), www.uniprot.org, April 14, 2023) in addition to the default MaxQuant contaminants. Searches were performed with a false discovery rate of 1%. A maximum of two missed tryptic cleavages was allowed. Searches were performed with a fixed modification for carbamidomethylation of cysteine residues and a variable modification for the oxidation of methionine and acetylation of the protein N-terminus. Label-free quantitation was performed using MaxLFQ with Fast LFQ, minimum ratio count of 2 and with match between runs [29]. Data analysis was performed using the Perseus computational platform for comprehensive analysis of proteomics data [30]. The mass spectrometry proteomics data have been deposited to the ProteomeXchange Consortium via the PRIDE partner repository with the dataset identifier PXD059011 [31].

### Mass spectrometry analysis of metabolites

Ultra-high performance liquid chromatography coupled with mass spectrometry (UHPLC/MS) analyses were conducted using a Thermo Scientific Horizon Flex UHPLC system, interfaced with a Thermo Scientific Orbitrap ID-X Mass Spectrometer. For the separation of polar metabolites, a HILICON iHILIC-(P) Classic HILIC column (100 x 2.1 mm, 5 µm) with a HILICON iHILIC-(P) Classic guard column (20 x 2.1 mm, 5 µm) was utilized. The mobile-phase solvents consisted of solvent A = 20 mM ammonium bicarbonate, 2.5 µM medronic acid, 0.1% ammonium hydroxide in 95:5 water:acetonitrile and solvent B = 2.5 µM medronic acid in 95:5 acetonitrile:water. The column compartment temperature was maintained at 45°C, and metabolites were eluted using a linear gradient at a flow rate of 0.25 mL/min as follows: 0-1 min, 90% B; 12 min, 35% B; 12.5-14.5 min, 25% B; 15 min, back to 90% B. The injection volume was 4 µL. Data were collected with the following settings: spray voltage, 3.5 kV (positive) and -3 kV (negative); sheath gas, 35; auxiliary gas, 10; sweep gas, 1; ion transfer tube temperature, 250°C; vaporizer temperature, 300°C; mass range, 70 – 1000 Da; resolution, 120,000 (MS1), 30,000 (MS/MS); collision energy, 30; maximum injection time, 100ms; isolation window, 1.6 Da. The LC/MS data were then processed and analyzed using Skyline [32].

### Quantification and statistical analysis

GraphPad Prism 9 was used for statistical analysis, specifically employing analysis of variance (ANOVA). The number of biological replicates for each experiment along with all statistical parameters, are detailed in the figure legends of graphs and tables. For clarity, only relevant statistical comparisons are shown in the graphs. Unless otherwise indicated, each data point represents the mean ± SD (n ≥ 3).

#### Results

### Pyrimethamine suppresses NRF2 mRNA and protein independently of KEAP1

PYR was discovered as an NRF2 inhibitor in a high-throughput small molecule screen of H1299- NQO1-eYFP lung cancer cells [24]. These cells constitutively express nuclear-localized mCherry and harbor an eYFP cassette stably integrated into the first intron of the *NQO1* gene, a canonical NRF2 target gene. To confirm PYR action and define its dose-response relationship, H1299- NQO1-eYFP cells were treated with the NRF2 inducing agents: CDDOme, sulforaphane or PRL295 with increasing amounts of PYR. Mechanistically, CDDOme and sulforaphane covalently modify and inactivate KEAP1[3]. PRL295 non-covalently binds KEAP1 to sterically occlude NRF2 binding[33]. PYR suppressed *NQO1*-driven eYFP induced by each NRF2 agonist, with an average IC50 of 1.2µM (**Figure 1A**). Western blot analysis of KYSE70 cells, which harbor an activating mutation in NRF2, confirmed PYR suppression of NRF2 protein levels and NRF2 target genes SLC7A11, HMOX1, and GCLC (**Figure 1B**). PYR also induced DHFR protein levels, which is known to tightly correlate with the inhibition of DHFR enzymatic activity[34]. In a time course analysis, PYR significantly decreased NRF2 and its target genes SLC7A11, HMOX1, and GCLC at 48h post-treatment; statistically insignificant suppression was observed at 24-36h (**Figure 1C**). To evaluate cell type specificity, a panel of NRF2-active cell lines was treated with PYR before w.blot analysis of NRF2, NRF2-target genes and DHFR (see **Table S1** for cell line genotypes). With varying efficacy, PYR decreased NRF2 protein abundance and NRF2 target gene expression, and increased DHFR protein levels in all cell types tested (**Figure 1D**).

**Figure 1.**
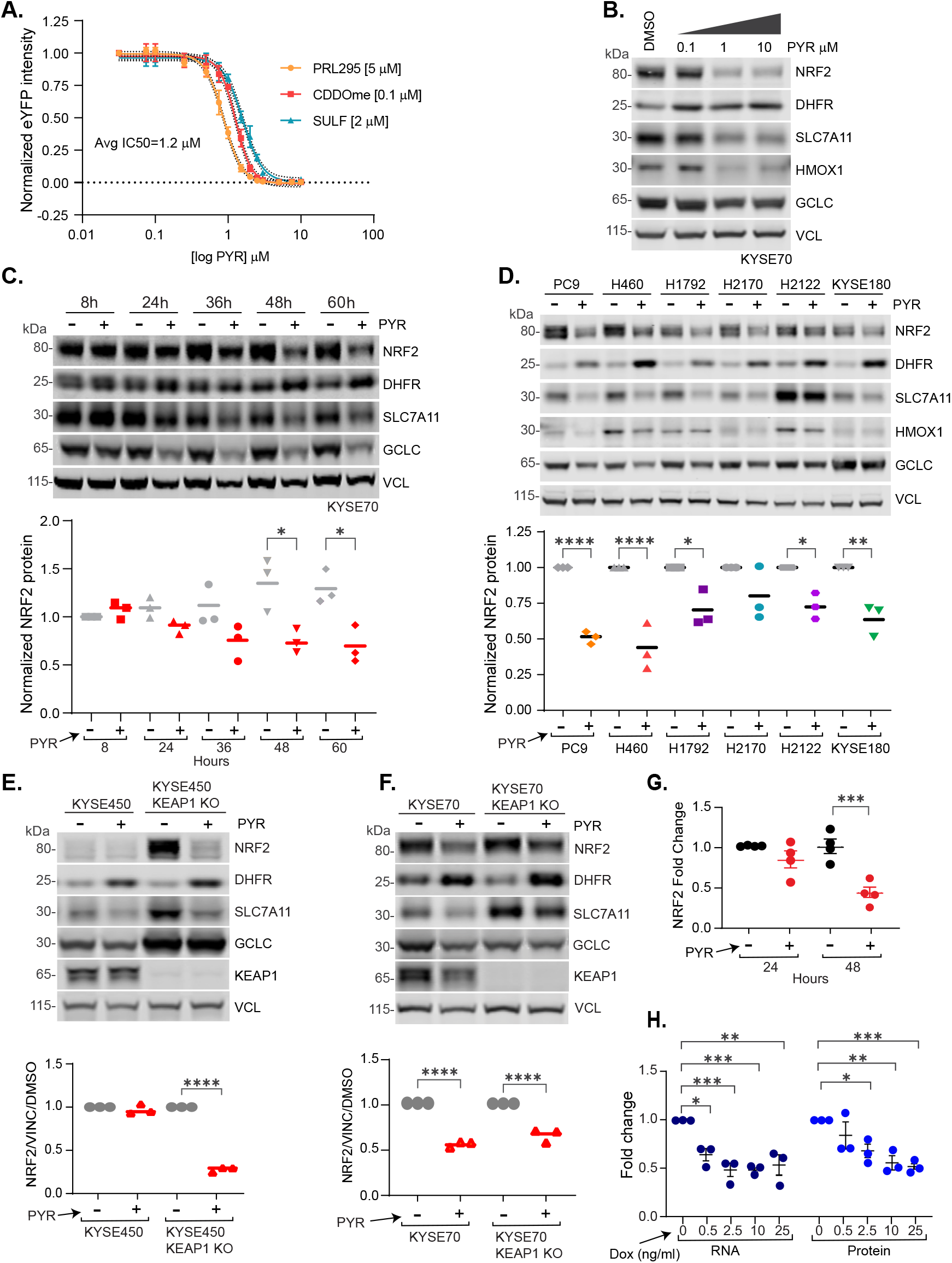
Pyrimethamine inhibits NRF2 mRNA and protein independently of KEAP1. **A.** H1299 NQO1-eYFP cells were treated with DMSO vehicle or 10 µM PYR in the presence of PRL295, CDDOme, or sulforaphane. **B.** Western blot analysis showing dose-dependent effects of PYR on indicated proteins in KYSE70 cells. **C.** Western blot analysis of PYR (10 µM) time course effects on indicated proteins in KYSE70 cells. Quantitation across biological triplicate experiments is shown below, normalized to VCL (vinculin) (*P < 0.05 by one-way ANOVA). **D.** Western blot analysis of lung and esophageal cancer cells following 48h treatment with PYR (10µM). Quantitation across biological triplicate experiments is shown below, normalized to VCL (*P < 0.05, **P < 0.01, ***P < 0.001, and ****P < 0.0001 by one-way ANOVA). **E-F.** Western blots assessing the effects of PYR (10µM; 48h) on the indicated proteins in KYSE70, KYSE450, or KEAP1 KO derivatives. Quantitation across biological triplicate experiments is shown below (****P < 0.0001 by one-way ANOVA). **G.** Graph showing *NRF2* mRNA fold change in KYSE70 cells following 24 and 48h treatment with PYR (10µM). NRF2 expression was normalized to *RPL13*A. Data are presented as mean ± SD (*P < 0.05, **P < 0.01, ***P < 0.001) by one-way ANOVA (n ≥ 3 biological replicates per group). **H.** KYSE70 cells harboring a dox-inducible NRF2 shRNA were treated with the indicated amount of dox for 24h before qPCR and western blot analysis for NRF2. NRF2 protein was normalized to VCL; *NRF2* mRNA expression was normalized to RPL13A. Data are presented as means ± SD (*P < 0.05, **P < 0.01, ***P < 0.001) by one-way ANOVA (n ≥ 3 biological replicates per group). See also Figure S1.

Because the KEAP1/CUL3 E3 ubiquitin ligase complex is a major cellular mechanism responsible for NRF2 degradation, we tested if KEAP1 was required NRF2 suppression by PYR. KYSE450 esophageal cells contain a wildtype KEAP1/NRF2/CUL3 pathway, and consequently low baseline NRF2 activity. CRISPR KO of KEAP1 induced NRF2 and NRF2 target gene protein expression (**Figure 1E**). PYR robustly suppressed NRF2 and SLC7A11 protein levels, but not GCLC. Similar results were seen in KEAP1 KO KYSE70 cells (**Figure 1F**). In both cell models, the presence of KEAP1 did not impact DHFR stabilization following PYR treatment. These KEAP1 KO data complement our findings that PYR inhibits NRF2 activation induced by KEAP1 chemical inhibitors (eg. CDDOme, Sulforaphane), and strongly suggests that PYR suppresses NRF2 independently of KEAP1. A simple mechanism for PYR-induced NRF2 suppression could be downregulation of NRF2 mRNA abundance. We analyzed NRF2 mRNA expression using qPCR following treatment of KYSE70 cells with PYR for 24h and 48h. While a modest decrease in *NRF2* mRNA expression was observed after 24h PYR treatment, *NRF2* mRNA expression decreased by 50% at 48h (**Figure 1G**). To determine if a 50% reduction in *NRF2* mRNA would impact NRF2 protein levels, we engineered KYSE70 cells with doxycycline-inducible short hairpin RNA (shRNA) targeting *NRF2* or luciferase as a control. Increasing doses of doxycycline dose-dependently suppressed both NRF2 protein and mRNA (**Figure 1H and S1**). These data suggest that PYR-induced NRF2 suppression may occur at the level of NRF2 mRNA expression.

### Structure-activity relationships across a series of pyrimethamine analogs

DHFR inhibition by methotrexate clinically benefits patients suffering many different cancer types. However, cancer-appropriate dosages of methotrexate carry severe toxicities to normal tissues, limiting its therapeutic potential. We and others have sought new drugs to inhibit DHFR and one-carbon metabolism[22]. To improve the potency of PYR and reveal insights into its mechanism of NRF2 suppression, we designed, synthesized, and tested 24 PYR analogs across two rounds of structure-activity relationship (SAR) profiling (see supplemental methods and **Figure S3**). Analogues #101 through #113 were based on PYR and removed and or modified the: 1) 4-amino moiety, 2) 6-ethyl moiety, or 3) the chlorine on the 5-*para*-chlorobenzene ring (**Figure 2A** and **Table S2**). All analogs were independently tested for suppression of NRF2-dependent transcription in H1299-NQO1-eYFP cells treated with CDDOme, sulforaphane, or PRL295. Analogs that suppressed eYFP expression at 10μM were further studied across a 12-point dose-response curve. Eliminating the 4-amino and 6-ethyl moieties abolished NRF2 suppression (#101 and #103, respectively). In contrast, the removal of the chlorine on the 5-*para*-chlorobenzene ring in #111 did not impact the potency of NRF2 suppression. Various substitutions of this chlorine atom eliminated NRF2 suppressive activity. However, moving the chlorine to the 3^rd^ position of the benzene ring strongly increased the potency of NRF2 suppression, with PYR demonstrating a 1.23µM IC50 compared to WCDD104 at an IC50 of 0.098µM (**Figure 2A** and **S2A-C, Table S2**). Analogs #114 through #134 considered additional modifications to WCDD104. Of these, WCDD114 and WCDD115 demonstrated the strongest potency for NRF2 suppression at 0.108 and 0.057µM, respectively (**Figure 2B-D**). To test if WCDD104, WCDD114, and WCDD115 suppress endogenous metrics of NRF2 signal transduction, we performed quantitative Western blot analysis on treated KYSE70, A549, OE21, and PC-9 cells, all of which harbor activating mutations in the NRF2 pathway (**Table S1**). DMSO and WCDD101, which had no impact on NRF2-driven NQO1-eYFP expression in H1299 cells, were used as negative controls while PYR served as a positive control. Treatment with PYR and each analog suppressed NRF2 protein levels and the expression of SLC7A11, HMOX1, GCLC, and NQO1 (**Figure 2E-H and S2D**). KEAP1 dependency was also evaluated for these PYR analogues, using parental and KEAP1 KO KYSE70 and KYSE450 cells. As expected, PYR analogues suppressed NRF2 irrespective of KEAP1 expression (**Figure S2 E, F**).

**Figure 2.**
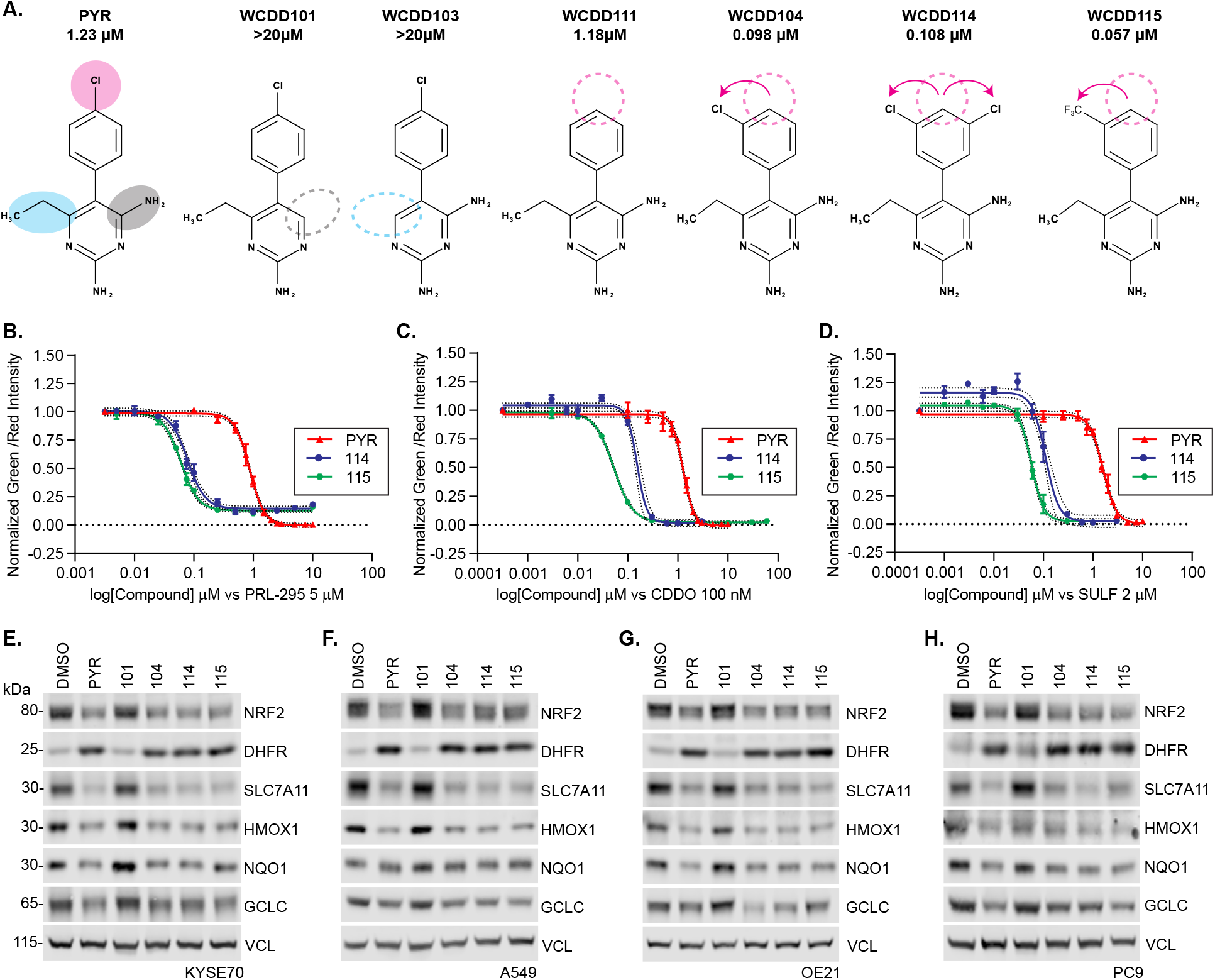
Structure-activity relationships across 25 pyrimethamine analogs. **A.** Chemical structures and IC50 values of PYR and select structural analogs. IC50 values reflect suppression of eYFP quantitation in H1299 NQO1-eYFP cells (see also Figure S3 and Table S1). **B-D.** Dose- response relationships for PYR, WCDD114, and WCDD115 in H1299 NQO1-eYFP cells treated with 5µM PRL295 (B), 100nM CDDOme (C), and 2µM SULF (D). eYFP intensity was normalized to mCherry. **E-H.** Western blots assessing the effects of 48h treatments with 10 µM PYR, 1 µM of WCDD114, WCDD115, WCDD101, and WCDD104 on KYSE70, A549, OE21, and PC9 cells. Quantitation and statistical analysis of biological triplicate experiments is shown in Figure S2D.

### DHFR chemical inhibition suppresses NRF2

Given that PYR inhibits hDHFR enzymatic activity, we next tested if DHFR inhibitors unrelated to PYR also suppressed NRF2. H1299-NQO1-eYFP cells were treated with increasing doses of methotrexate (MTX), cycloguanil (CG), or pemetrexed (PEM) in the presence of the NRF2 activating compounds PRL295, CDDOme, or SULF (**Figure 3A-C**). MTX and CG efficaciously inhibited *NQO1* driven eYPF expression, as did PYR. PEM, a multitarget antifolate known to inhibit DHFR, thymidylate synthase, and glycinamide ribonucleotide formyl transferase, suppressed eYFP expression but with much weaker efficacy. Western blot analysis of NRF2- active KYSE70 and A549 cells confirmed that PYR, WCDD115, MTX, and CG suppressed endogenous NRF2 signaling and increased DHFR protein (**Figure 3D, E and S4**). In contrast, PEM did not inhibit NRF2 and weakly increased DHFR protein levels. At the RNA level, MTX, PYR, and WCDD115 suppressed the expression of *NRF2*, *NQO1*, *GCLC*, and *SXRN1* with similar efficacy (**Figure 3F** and **S4**). To directly establish the novel WCDD115 compound as an hDHFR inhibitor, we performed an *in vitro* enzyme activity test of recombinant hDHFR (**Figure 3G**). Across 6 biological replicate experiments, the IC50 for hDHFR inhibition was: 0.006µM for MTX, 4.49µM for PYR and 0.144µM for WCDD115, establishing that WCDD115 is 31-fold more potent that PYR in suppressing hDHFR. Together, these data suggest that WCDD115 and structurally distinct DHFR inhibitors suppress NRF2 expression and its downstream target genes.

**Figure 3.**
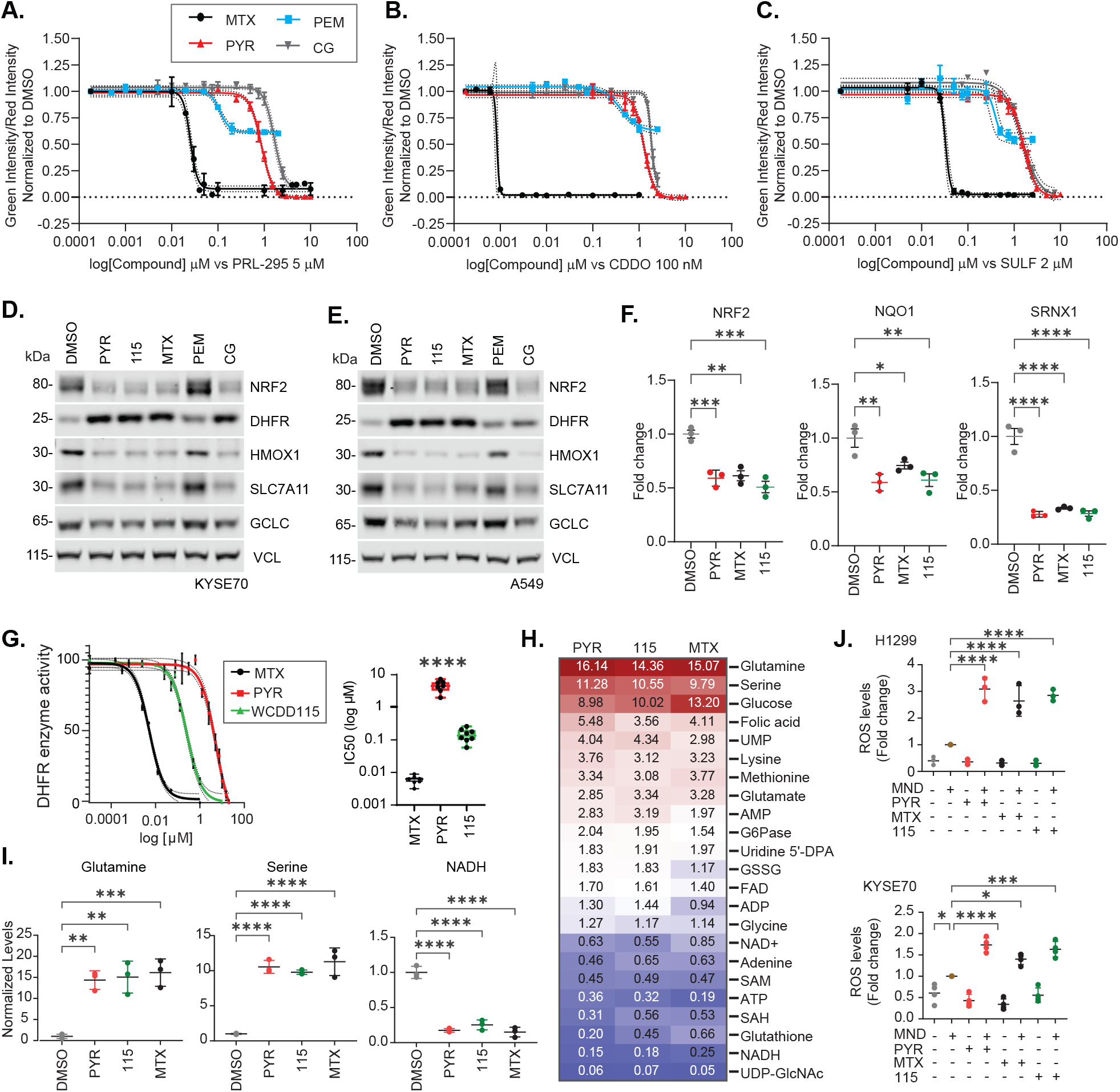
WCDD115 and chemically distinct DHFR inhibitors suppress NRF2 and reprogram cellular metabolism. **A-C.** H1299 NQO1-eYFP cells were treated with increasing doses of the indicated DHFR inhibitors in the presence of PRL295, CDDOme, or SULF. eYFP intensity was normalized to cell number (mCherry positive) and DMSO vehicle. **D, E.** Western blot analysis of KYSE70 and A549 cells treated with PYR (10µM), WCDD115 (1µM), MTX (0.1µM), PEM (100nM), or CG (10μM) for 48h. Quantitation and statistical analysis is shown in Figure S4. **F.** *NRF2* mRNA was quantified by qPCR after 48h treatment of KYSE70 cells with PYR (10µM), WCDD115 (1µM), MTX (0.1µM). Data are normalized to RPL13A and plotted as mean ± SD (*P < 0.05, **P < 0.01, ***P < 0.001) by one-way ANOVA. **G.** Dose-response relationships for PYR, WCDD115, and MTX on the enzymatic activity of recombinant hDHFR (3E-3 units). Data are normalized to vehicle and presented as mean ± SD (n=7, ****P< 0.001) by one-way ANOVA. **H.** Targeted metabolomic profiles of KYSE70 cells treated with PYR (10µM), WCDD115 (1µM), MTX (0.1µM) for 48h. Data shown are fold changes from DMSO (see also Table S3). **I.** Glutamine, Serine, and NADH levels following 48h treatment of KYSE70 cells with PYR (10µM), WCDD115 (1µM), and MTX (0.1µM). Data are presented as means ± SD (*P < 0.05, **P < 0.01, ***P < 0.001) by one-way ANOVA. **J.** H1299 and KYSE70 cells were treated with PYR (10µM), WCDD115 (1µM), and MTX (0.1µM) 48 h ± 20µM menadione for the last 3h. ROS was quantified using CellRox. Data are presented as means ± SD (*P < 0.05, **P < 0.01, ***P < 0.001, ****P< 0.001) by one-way ANOVA (n≥3 biological replicates per group).

DHFR is a critically important enzyme within the one-carbon metabolism circuitry. Its chemical or genetic loss results in well-characterized alterations to the global metabolome, including specific amino acids, purines, pyrimidines, methylation events, NADH and global redox status[35]. We used targeted mass spectrometry to quantify 48 metabolites following PYR, WCDD115, or MTX treatment in KYSE70 and H1299 cells (**Figure 3H** and **Table S3**). Overall, highly similar metabolic profiles were induced by all three DHFR inhibitors. The identities of the impacted metabolites were consistent with the known roles of DHFR in one-carbon metabolism. For example, each compound resulted in an accumulation of folic acid, glucose, glutamine, and serine with decreases in NADH and glutathione levels (**Figure 3H, I**).

Both NRF2 transcriptional activity and one-carbon metabolism facilitate cellular redox balance, mostly through the production of glutathione, NADH and NADPH[36, 37]. To determine if DHFR inhibition with WCDD115, PYR, or MTX suppresses cellular capacity to neutralize ROS, we treated KYSE70 cells and H1299 cells in the presence and absence of a low, sensitizing dose of menadione (MND) to increase endogenous ROS. In the absence of MND, the compounds had no significant effect on the CellRox ROS-sensitive dye (**Figure 3J**). However, in MND-treated cells, PYR, WCDD115, and MTX induced ROS levels in both cell models. Thus, the chemical suppression of DHFR by PYR, its more potent analog WCDD115, and the structurally dissimilar MTX compound, suppress NRF2, yield similar metabolic profiles consistent with loss of one-carbon metabolism, and suppress the cellular capacity to neutralize ROS.

### Inhibition of DHFR is required for WCDD115-mediated suppression of NRF2

DHFR reduces dietary folic acid to tetrahydrofolate (THF), which then accepts one-carbon units (methyl groups) from serine to form precursors for purine synthesis, thymidylate synthesis, and the methionine cycle. Seeking to determine if DHFR inhibition was required for NRF2 suppression, we performed two metabolic rescue experiments. First, folinic acid (FA) is a THF derivative that does not require DHFR reduction to initiate one-carbon transfer reactions; it replenishes the pathway epistatically downstream of DHFR. KYSE70 cells were treated with PYR, MTX, WCDD115, PEM, or CG alone or in combination with FA before Western blot analysis of NRF2, DHFR, and the NRF2 target genes HMOX1 and SLC7A11 (**Figure 4A**). For each DHFR inhibitor including WCDD115, FA co-treatment restored NRF2 expression. Second, we tested if supplementation of exogenous hypoxanthine (H) and thymidine (T) rescued WCDD115-mediated suppression of NRF2. H and T feed the purine and pyrimidine synthesis pathways downstream of DHFR and THF. Like FA, HT treatment blocked NRF2 suppression by WCDD115 in KYSE70 cells (compare lanes 2 & 4, **Figure 4B**). HT treatment alone did not impact NRF2 protein levels (lanes 1 & 3, **Figure 4B**).

**Figure 4.**
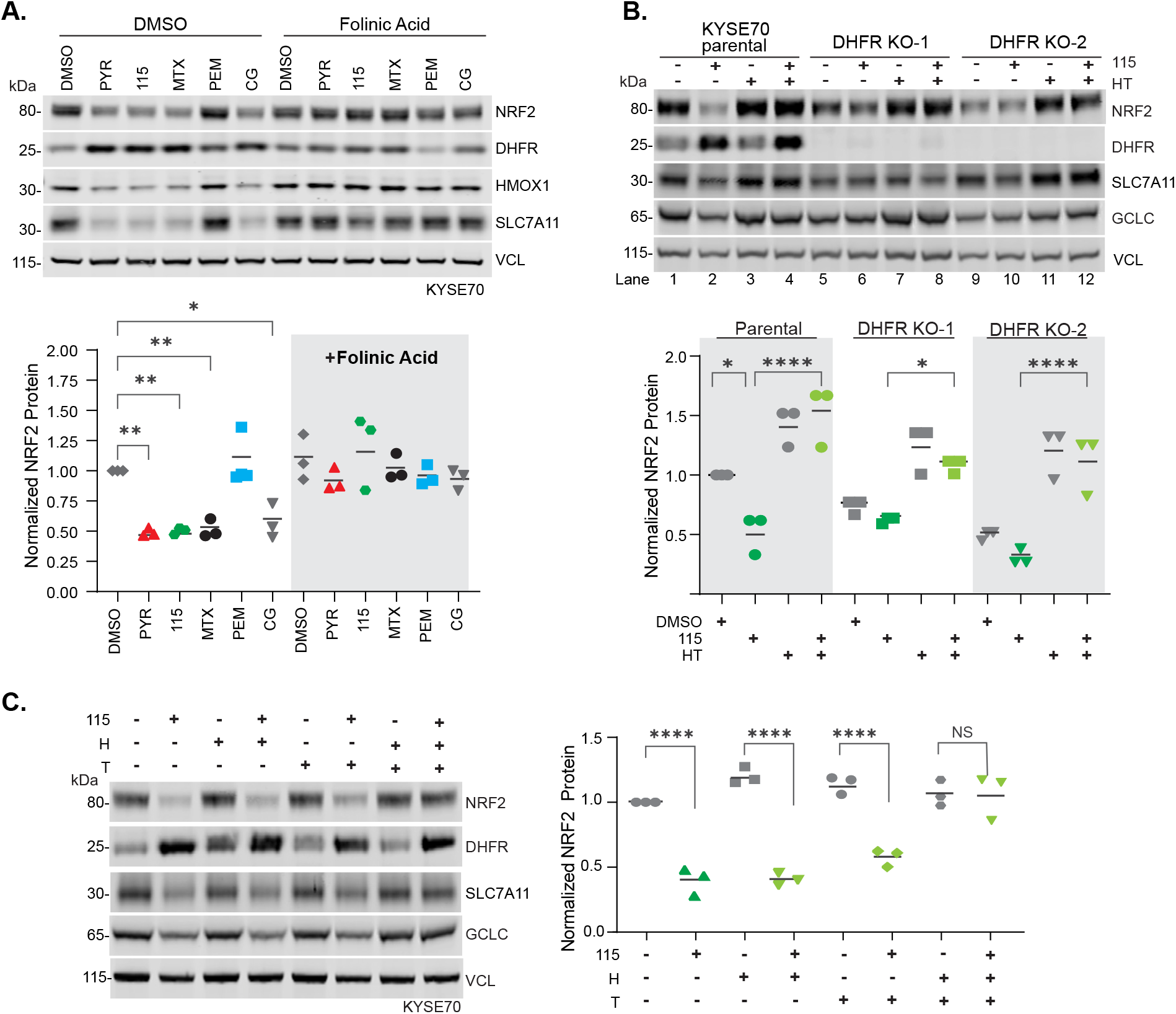
Inhibition of DHFR is required for WCDD115-mediated suppression of NRF2. A. Western blot analysis of KYSE70 cells treated with PYR (10 µM), WCDD115 (1 µM), MTX (0.1 µM), PEM (0.1 µM), or CG (10 µM) in the absence or presence of folinic acid (10mg/ml). Data are presented as means ± SD (*P < 0.05, **P < 0.01) by one-way ANOVA (n=3 biological replicates per group). **B.** KYSE70 parental cells or two monoclonal DHFR KO derivatives were treated with WCDD115, HT, or both for 48 h. Protein expression of DHFR, NRF2, and downstream targets SLC7A11, HMOX1, NQO1, GCLC, and VINC was quantified by Western blot. Quantitative data of biological triplicate experiments are plotted below as mean ± SD (*P < 0.05, **P < 0.01, ***P < 0.001, ****P < 0.001) by one-way ANOVA. **C.** KYSE70 cells were treated with the indicated combinations of WCDD115, hypoxanthine (H), and thymidine (T) for 48h. Protein expression of DHFR, NRF2, and downstream targets SLC7A11, HMOX1, NQO1, GCLC, and VINC was quantified by Western blot. Quantitative data of biological triplicate experiments are plotted below as mean ± SD (*P < 0.05, **P < 0.01, ***P < 0.001, ****P < 0.001) by one-way ANOVA.

We next tested if genetic knockout of DHFR phenocopied chemical inhibition with respect to NRF2 suppression. As an essential gene, the viability of DHFR KO cells requires HT supplementation (or FA). We used HT-containing media and CRISPR-Cas9 to create two monoclonal DHFR KO KYSE70 cell lines. HT removal from each DHFR KO cell line resulted in suppression of NRF2 and NRF2 target genes (lane 1 vs. 5 & 9, **Figure 4B**). Importantly, in the DHFR KO cells cultured without HT, WCDD115 had no effect on NRF2 protein levels (lane 5 vs. 6 and 9 vs. 10, **Figure 4B**). These data indicate that with respect to NRF2, DHFR genetic loss phenocopies DHFR chemical suppression and that WCDD115 requires DHFR to inhibit NRF2 protein expression. Last, we determined whether purines (H) or pyrimidines (T) differentially rescued NRF2 suppression by WCDD115. Western blot analysis of KYSE70 cells treated with H, T, or HT in the presence and absence of WCDD115, shows that both H and T are required to metabolically rescue WCDD115-medaited suppression of NRF2 (**Figure 4C**).

### WCDD115 is an indirect inhibitor of NRF2

Pyrimethamine is reported to inhibit STAT3 signaling, and our recent report and this work shows that it also inhibits NRF2 [19, 20]. Additional studies connect PYR to suppression of NF-kB, p38, and DX2 as well as epithelial to mesenchymal transition and cancer cell invasion, among other biologies [38, 39]. Therefore, we used mass spectrometry-based proteomics to evaluate the global impact of PYR, WCDD115, and MTX on the proteome. In a targeted approach, an optimized internal standard triggered-parallel reaction monitoring (OIS-PRM) method was used to quantify a predefined panel of proteins [26]. This panel contains 158 peptides mapping to 70 proteins, which encompass NRF2, NRF2 target genes, KEAP1, DHFR, and cell cycle proteins. Treatment of KYSE70 cells with WCDD115 significantly suppressed 29 proteins, including NRF2 and 20 NRF2 targets (**Figure 5A**, **S5A, Table S4**). Comparison of WCDD115 to PYR and MTX revealed remarkable overlap (**Figure 5B**). DHFR, TALDO1, and UGDH were among the most increased proteins. Next, we complemented the targeted MS with unbiased, data-dependent acquisition mass spectrometry (DDA-MS). KYSE70 cells were treated with WCDD115, PYR, or MTX before DDA-MS, revealing 4673 quantified protein groups. Of these, ∼2400 proteins were statistically increased or decreased in abundance, with a strong overlap of 2,194 proteins between the three compounds (**Figure 5C**, **Table S5**). Interestingly, the polarity of affected proteins was balanced (**Figure 5D,E**, **S5B**). These results suggest that DHFR inhibition by WCDD115, PYR, and MTX results in similar impacts on the proteome characterized by many hundreds of proteins that increase and decrease. Gene set enrichment analysis on the differentially expressed proteins revealed upregulation of DNA repair, E2F targets, p53, TNFα, and interferon-gamma ontologies. Down-regulated enrichments included NRF2-associated ROS signaling, the PI3K/AKT/MTOR pathway, and cholesterol homeostasis (**Figure 5F**, **Table S6**).

**Figure 5.**
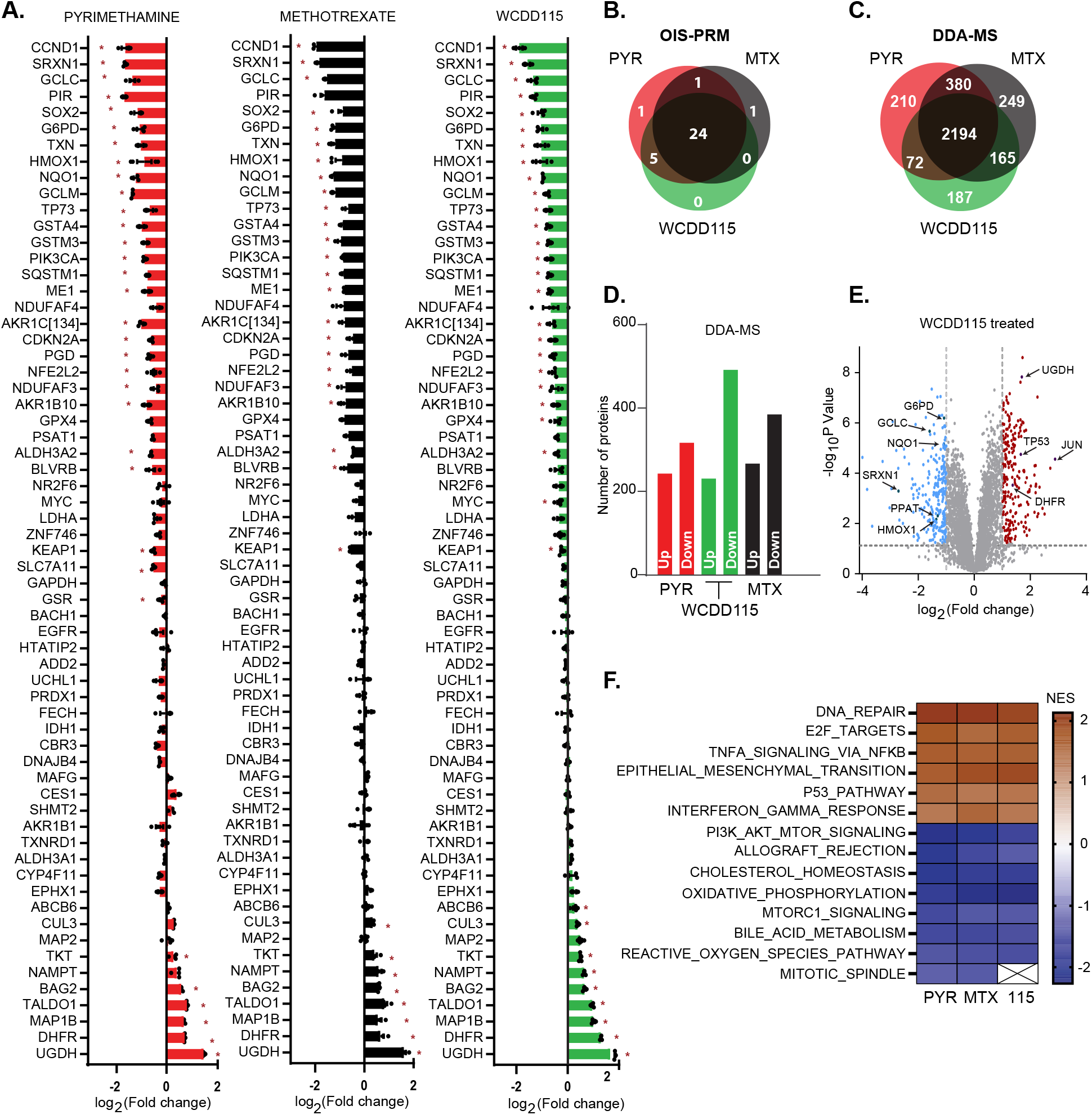
WCDD115 is an indirect inhibitor of NRF2. **A.** KYSE70 cells were treated with PYR (10 µM), WCDD115 (1 µM), and MTX (0.1 µM) for 48h before OIS-PRM analysis. Data are presented as log2-transformed geometric mean of the peak area ratios for peptides mapping to the indicated protein. Significance was calculated using two-way ANOVA and Šidák test for multiple comparisons (*P <0.05). Data represent mean ± SD across biological triplicate experiments. **B.** Venn diagram of OIS-PRM results shown in (A) reflecting the number of proteins differentially induced or repressed >2-fold. **C.** Protein samples shown in (A) were analyzed via data-dependent acquisition (DDA). Proteins increased or decreased by 2-fold or greater are comparatively depicted by a Venn diagram. **D.** Histogram plot showing the number of upregulated and downregulated proteins by 2-fold. **E.** Volcano plot visualization of protein changes following WCDD115 treatment of KYSE70 cells. Blue and red data points represent significantly (p< 0.05) downregulated and upregulated proteins, respectively (log2 fold-change ≥1) (see also Fig. S5B). **F.** Partial list of Hallmark terms identified through gene set enrichment analysis (GSEA) of differentially expressed proteins (see also Table S5).

### Discussion

The NRF2 transcription factor and oncogene is an important and promising drug target in oncology. Most cancers harboring activating mutations within the NRF2 pathway are addicted to NRF2, and thus therapeutic inhibitors of NRF2 may show single-agent efficacy[40, 41]. That said, in all cancers irrespective of NRF2 pathway addiction, NRF2-targeting agents may increase the efficacy of front-line chemotherapy, radiation therapy, and immune checkpoint inhibitors. Such predictions are well-supported across decades of pre-clinical genetic and pharmacological studies. FDA-approved NRF2 inhibitors are currently lacking. In academia and industry, drug development programs have identified many NRF2 inhibitors that show cell-based and/or animal model activity, including ML385, brusatol, halofuginone, MSU38225, and VVD-8065 [16, 42]. We contributed to this effort with the identification of PYR and 11 lead compounds from a chemical screen of NRF2-dependent transcription in H1299 cells [19]. Here we report new insights into PYR as an NRF2 inhibitor, including its mechanism of action via suppression of 1C metabolism and the development of WCDD115 as a 22-fold more potent PYR analogue.

A primary conclusion drawn from this study is that 1C metabolism is required for NRF2 pathway activity, but the precise mechanism that connects 1C metabolism to NRF2 remains unresolved. Genetic knockout of DHFR and three structurally distinct chemical inhibitors of DHFR decrease NRF2 mRNA and protein levels. These data support a simple model that following suppression of 1C metabolism, loss of NRF2 mRNA results in loss of NRF2 protein. To test this, future experiments should engineer NRF2 mRNA levels to remain high despite DHFR inhibition, perhaps through CRISPR-A or transgene approaches. Metabolite rescue experiments with hypoxanthine and thymidine supplementation indicate that NRF2 expression is particularly sensitive to purine and thymidine pools generated by 1C metabolism. Testing whether DHFR inhibition and/or loss of nucleotide pools impacts the activity of the NRF2 promoter is needed, particularly in light of recent studies suggesting that DHFR may act as an enzymatically active transcriptional co- activator of STAT3-mediated transcription [20]. This DHFR-centric model complements and contrasts our previous report showing that PYR induces KEAP1-independent NRF2 ubiquitylation, resulting in decreased NRF2 protein half-life [19]. It remains unknown if PYR and WCDD115 directly facilitate NRF2 ubiquitylation or if the cell responds to DHFR inhibition with increased NRF2 ubiquitination.

The global and targeted proteomics data revealed highly concordant responses to MTX, PYR and WCDD115, including roughly equivalent numbers of up-and down-regulated proteins that aligned well across gene ontological enrichments. Thus, DHFR chemical inhibition does not globally suppress transcription, translation, or proteostasis, but selectively increases or decreases a host of cellular signaling pathways and biologies. Whereas NRF2 and the PI3K/AKT/MTOR pathway are suppressed, DNA repair, NFκB, p53 and epithelial to mesenchymal transition (EMT) are induced. These data support and contrast recently published findings [20, 21]. First, we and others observe that nucleotide pool restrictions following DHFR inhibition induces the DNA damage and p53 response. Second, our proteomic data show that PYR/WCDD115/MTX increased the enrichment of EMT ontology in treated KYSE70 cells. In contrast, PYR but not MTX was recently shown to inhibit EMT and metastasis in lung cancer, potentially indicating cell-type specific responses to DHFR inhibition [21]. Last, suppression of 1C metabolism is known to result in an increase in reactive oxygen species, which we also observe. While suppressed NADPH and NADH production is likely key for ROS induction, it remains unknown if NRF2 loss by 1C suppression also contributes to ROS accumulation or ROS-driven cell toxicity.

PYR has received considerable attention in both drug repurposing efforts for oncology and as an alternative to methotrexate in rheumatological diseases [43–45]. The potential clinical value of PYR has spurred chemical explorations of the PYR backbone to improve its potency and efficacy; the discovery of WCDD115 contributes to these efforts. A recent study from the Page laboratory used SAR to identify two compounds with superior DHFR engagement in cells [46]. Consistent with our data, modifications to the 5-para-chlorobenzene ring proved most impactful for DHFR suppression. However, the Brown et. al. study concluded that the 6-ethyl moiety on PYR was not required for DHFR suppression, which directly contrasts our finding that WCDD103 lacking the 6- ethyl group has no effect on NRF2. Though we observe a strong correlation between the chemical inhibition of DHFR and NRF2, our SAR experiments did not directly test all PYR analogues on DHFR activity, but rather examined NRF2-mediated transcription (eg. WCDD103). Direct comparison of DHFR activity after PYR, WCDD115, and compounds 32, 34 of the Brown study is needed. Despite this, we suggest that WCDD115 represents a novel and potent inhibitor of human DHFR and may demonstrate improved anti-cancer efficacy as compared to PYR.

Several points of discussion highlight the clinical value of this work. Therapeutic targeting of 1C metabolism with MTX has been a cornerstone therapeutic approach in oncology and rheumatology. However, MTX carries significant dose-limiting toxicities, many of which have not been reported with PYR, including hepatotoxicity, GI intolerance, nephrotoxicity, and mucositis [47–50]. Further, MTX undergoes polyglutamation (PG) in cells to form MTXPG [51]. MTXPG levels correlate with activity and with adverse clinical events. Because PG activities vary across patients, toxicities and efficacy also fluctuate, limiting the clinical utility of MTX in oncology and rheumatology[52–54]. In contrast, PYR and WCDD115 do not undergo polyglutamation, thereby potentially improving its clinical efficacy. Finally, in contrast to MTX, PYR is comparatively safe during pregnancy. Pre-clinical head-to-head studies PYR, MTX and WCDD115 are needed to define their therapeutic value as MTX alternatives in cancer and rheumatology[45]. PYR was recently clinically evaluated in a phase I trial for chronic lymphocytic leukemia (CLL), proving it to be well-tolerated[23]. We are currently conducting an early phase I trial to evaluate the safety and biological activity of PYR in patients with locally advanced, HPV-unrelated head and neck squamous cell carcinoma (Clinical trial identifier: NCT05678348). Future clinical studies are needed to evaluate PYR as a combination agent chemotherapy or radiation therapy.

## Acknowledgments

We extend our gratitude to Dr. Dhaval Bhatt and Dr. Chase Weidmann for their insightful conversations and assistance with experimental troubleshooting. We also thank the members of the Major lab, Bernard Weissman lab, and Luke Chen lab for their expertise and support throughout this project. Finally, we thank and acknowledge Dr. Saira Sheikh and her team for their expertise in rheumatological diseases and the clinical rationale for MTX alternatives like PYR and WCDD115.

## Author Contributions

Conceptualization (JC, BMB, RD, PZ, MBM), Formal analysis (JC, BMB, EW, CP, KC); Funding acquisition (MBM, XC); Investigation (JC, BMB, KL, EW, MM, CP, KC); Supervision (GJP, RD, MBM); Roles/Writing - original draft (JC, MBM); and Writing - review & editing (XC, PZ, MBM).

## Data Availability Statement

All data presented in this study are available within the article, the supplementary data files, and within the PRIDE partner repository (PXD059011). All unprocessed, raw data are available upon request.

---

**Supplemental Figure 1.**
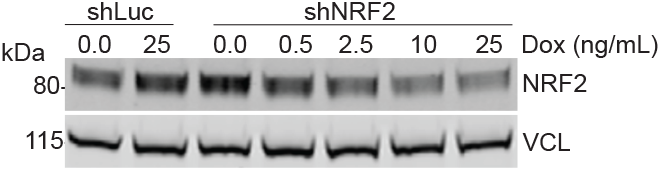
Correlations between *NRF2* RNA abundance and NRF2 protein abundance. Representative Western blot image used to produce quantitative data shown in Figure 1H.

**Supplemental Figure 2.**
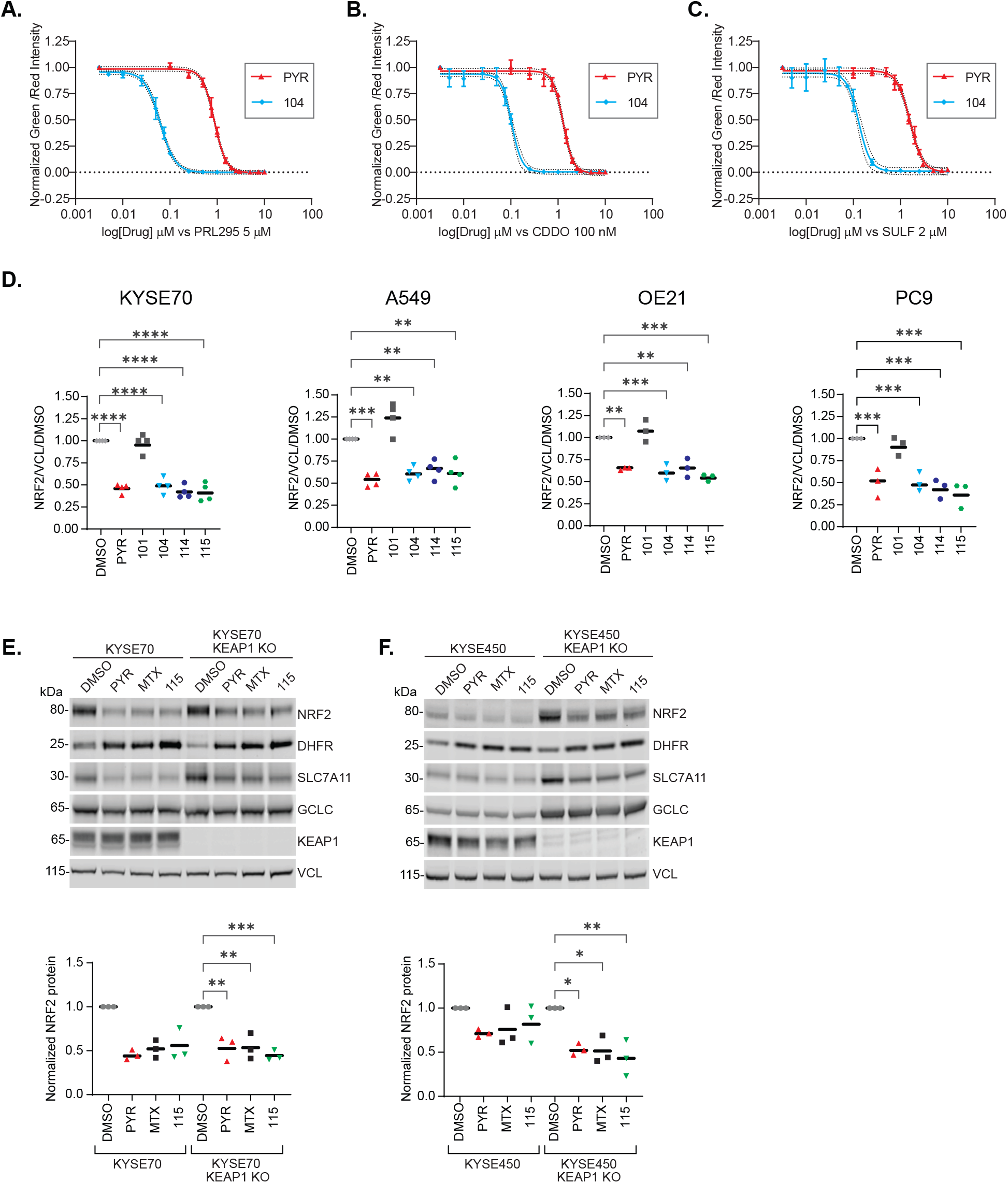
KEAP1-independent suppression of NRF2 by PYR and its structural analogues. **A-C.** H1299 NQO1-eYFP cells were treated with increasing doses of PYR or WCDD104 in the presence of PRL295, CDDOme or 2 µM SULF. eYFP intensity was normalized to cell number (mCherry positive) before normalization to DMSO vehicle. **D.** Quantified data from Western blots shown in Figure 2E-H (**P < 0.05, ***P < 0.01, ****P < 0.001,) by one-way ANOVA (n=4 biological replicates per group). **E-F.** Western blot analysis of KYSE70 and KYSE450 cells and their KEAP1 KO derivatives after 48h treatment with 10µM PYR, 1µM 115, or 0.1µM MTX. Below, quantitative, normalized data are plotted for NRF2/VCL.

**Supplemental Figure 3.**
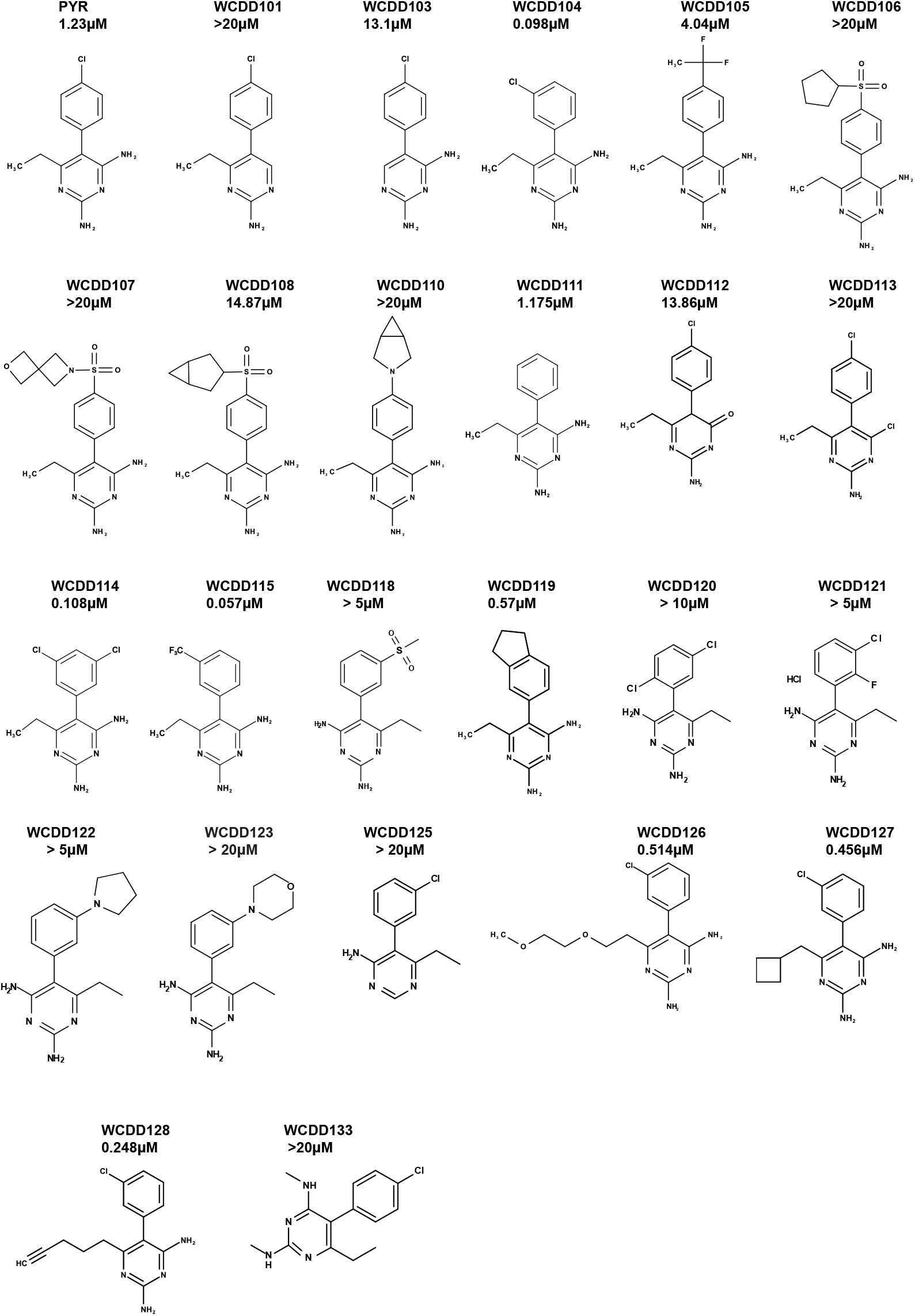
Structures and NRF2 inhibition IC50 values of PYR analogues. IC50 values were defined by the H1299-NQO1-eYFP assay, taking the average IC50 values from SULF, PRL295 and CDDOme (see Table S2).

**Supplemental Figure 4.**
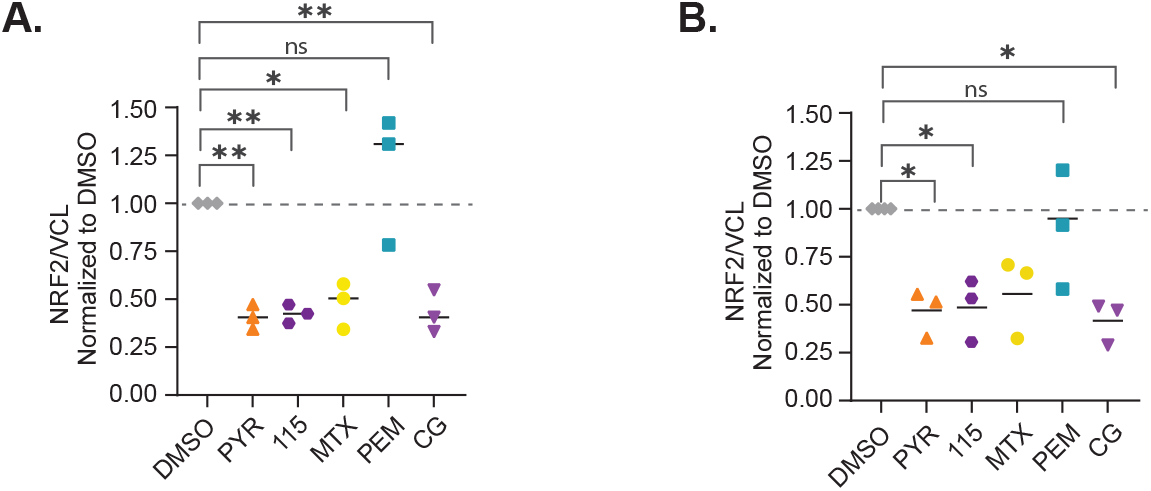
Chemically distinct DHFR inhibitors suppress NRF2 signaling. **A,B**. Quantitative Western blot data associated with Figure 3D and 3E. Protein expression was normalized to VCL and then DMSO (*P < 0.05, **P < 0.01, ***P < 0.001 by one-way ANOVA across biological triplicate experiments).

**Supplemental Figure 5.**
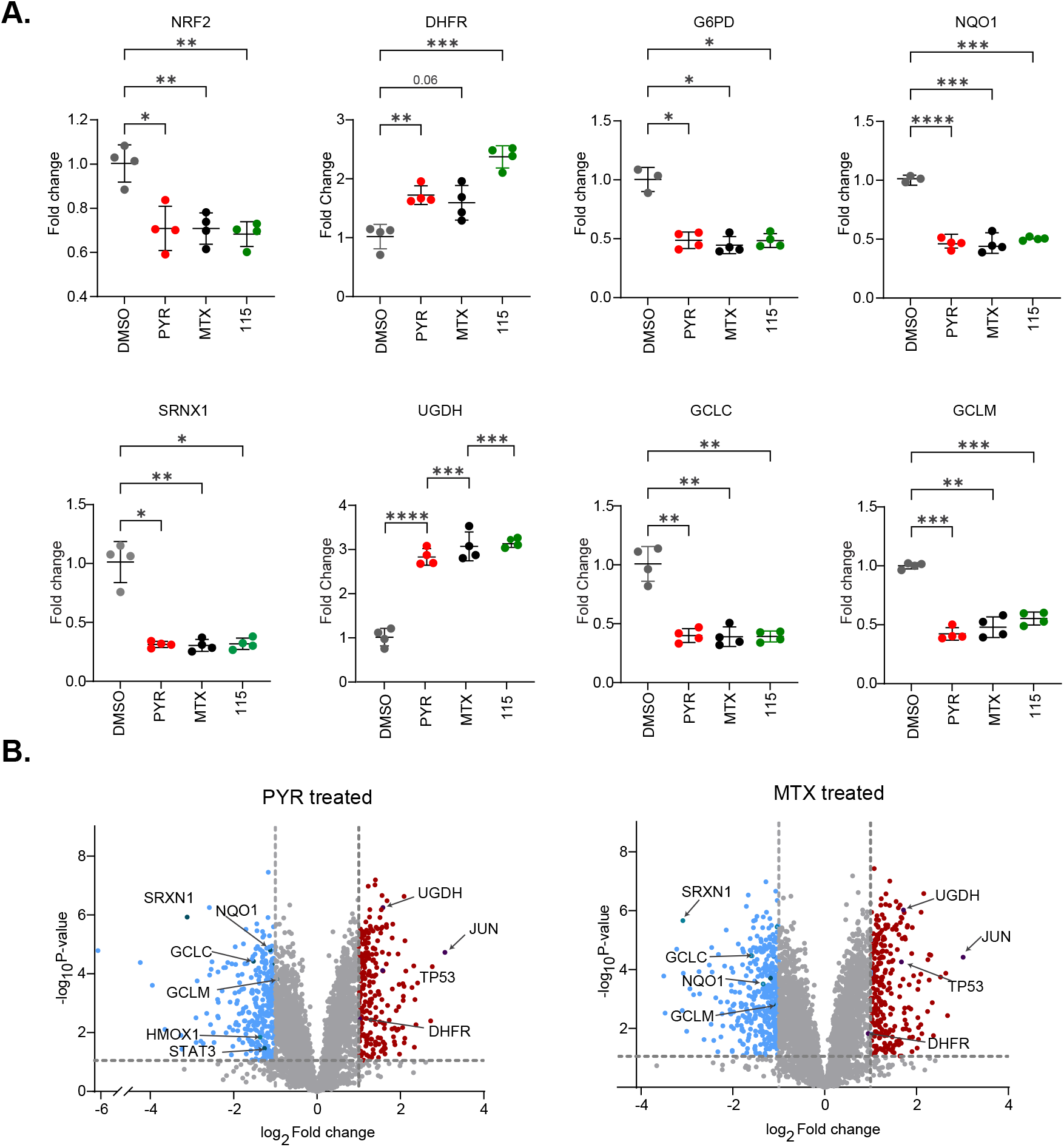
WCDD115 is an indirect inhibitor of NRF2. **A.** OIS-PRM derived quantitation of the indicated proteins, derived from Figure 5A. Horizontal lines denote mean values, and asterisks indicate levels of statistical significance (*p<0.05; **p<0.01; ***p<0.001; ****p<0.0001). **B**. Volcano plot showing differentially expressed proteins in KYSE70 cells after 48h treatment with PYR or MTX.

## Supplemental Tables

**Table S1.** Cell line mutation status for NFE2L2 (NRF2) and KEAP1. Mutation status for NFE2L2 (NRF2) and KEAP1 in cell line used in this study.

| Cell Line | NRF2 Alteration | KEAP1 Alteration |
| --- | --- | --- |
| KYSE70 | W24C |  |
| A549 |  | G333C |
| OE21 | G81S, D318H |  |
| PC9 | copy number amplification |  |
| H1299 |  |  |
| KYSE180 | D77V | P278Q |
| H460 |  | D236H |
| H2170 |  | R336* |
| H1792 |  | G462W |
| H2122 |  | A170_R204del |
| KYSE450 |  |  |

**Table S2.** Analog IC50 results in H1299-NQO1-YFP cells vs activators. Table showing the structures of PYR analogs generated in this study with IC50 values for NRF2 inhibition.

| Compound | CDDO 100nM | PRL295 5μM | SULF 2μM | Avg IC50 μM | Modified from |
| --- | --- | --- | --- | --- | --- |
|  | IC50 mM + SEM |  |  |  |  |
| PYR | 1.245 ± 0.024 | 0.873 ± 0.032 | 1.591 ± 0.0448 | 1.23 | N/A |
| WCDD101 | >20 | >20 | >20 | >20 | PYR |
| WCDD103 | 16.86 ± 2.563 | 9.99 ± 0.0737 | 12.46 ± 0.798 | 13.10 | PYR |
| WCDD104 | 0.103 ± 0.001 | 0.061 ± 0.003 | 0.129 ± 0.003 | 0.098 | PYR |
| WCDD105 | 5.115 ± 0.074 | 3.743 ± 0.4988 | 3.279 ± 0.006 | 4.04 | PYR |
| WCDD106 | >20 | >20 | >20 | >20 | PYR |
| WCDD107 | >20 | >20 | >20 | >20 | PYR |
| WCDD108 | 16.85 ± 0.208 | 15.21 ± 1.295 | 12.56 ± 0.1033 | 14.87 | PYR |
| WCDD110 | >20 | >20 | >20 | >20 | PYR |
| WCDD111 | 1.105 ± 0.016 | 1.511 ± 0.0329 | 0.910 ± 0.010 | 1.175 | PYR |
| WCDD112 | 19.84 ± 0.338 | 10.83 ± 0.620 | 10.92 ± 0.0655 | 13.86 | PYR |
| WCDD113 | >20 | >20 | >20 | >20 | PYR |
| WCDD114 | 0.134 ± 0.014 | 0.079 ± 0.002 | 0.111 ± 0.009 | 0.108 | WCDD104 |
| WCDD115 | 0.052 ± 0.0001 | 0.060 ± 0.001 | 0.057 ± 0.003 | 0.057 | WCDD104 |
| WCDD118 | >5 | >5 | >5 | >5 | WCDD104 |
| WCDD119 | 0.614 ± 0.021 | 0.552 ± 0.003 | 0.557 ± 0.03 | 0.57 | WCDD104 |
| WCDD120 | >10 | >10 | >10 | >10 | WCDD104 |
| WCDD121 | >5 | >5 | >5 | >5 | WCDD104 |
| WCDD122 | >5 | >5 | >5 | >5 | WCDD104 |
| WCDD123 | >20 | >20 | >20 | >20 | WCDD104 |
| WCDD125 | >20 | >20 | >20 | >20 | WCDD104 |
| WCDD126 | 0.461 ± 0.017 | 0.683 ± 0.002 | 0.397 ± 0.011 | 0.514 | WCDD104 |
| WCDD127 | 0.469 ± 0.002 | 0.444 ± 0.002 | 0.454 ± 0.006 | 0.456 | WCDD104 |
| WCDD128 | 0.244 ± 0.006 | 0.334 ± 0.004 | 0.163 ± 0.002 | 0.248 | WCDD104 |
| WCDD133 | >20 | >20 | >20 | >20 | WCDD104 |

**Table S3. Targeted metabolomic profiles of DHFR inhibitors.**

Table showing LC/MS metabolomics data following a 48h treatment of KYSE70 cells with DMSO, PYR, WCDD115, and MTX.

**Table S4. Targeted proteomic OIS-PRM data.**

Table showing peptide and protein log transformed abundances following a 48h treatment of KYSE70 cells with DMSO, PYR, WCDD115, and MTX. Data processing were performed as described in Wamsley, Nathan T. et al. Molecular & Cellular Proteomics, Volume 22, Issue 11, 100647.

**Table S5. Discovery DDA proteomic data.**

Table showing data-dependent acquisition proteomics data comparing treatment KYSE70 cells with DMSO, PYR, WCDD115, and MTX.

**Table S6. GSEA data derived from the DDA proteomics.**

Output table showing gene set enrichment analysis (GSEA) run against the Hallmark gene set.

## References

1. McMahon, M., et al., The Cap’n’Collar basic leucine zipper transcription factor Nrf2 (NF- E2 p45-related factor 2) controls both constitutive and inducible expression of intestinal detoxification and glutathione biosynthetic enzymes. Cancer Res, 2001. 61(8): p. 3299–307.

2. Itoh, K., et al., An Nrf2/small Maf heterodimer mediates the induction of phase II detoxifying enzyme genes through antioxidant response elements. Biochem Biophys Res Commun, 1997. 236(2): p. 313–22.

3. 3. Baird, L. and M. Yamamoto, The Molecular Mechanisms Regulating the KEAP1-NRF2 Pathway. 2020, Taylor & Francis.

4. Adinolfi, S., et al., The KEAP1-NRF2 pathway: Targets for therapy and role in cancer. Redox Biol, 2023. 63: p. 102726.

5. Yamamoto, M., T.W. Kensler, and H. Motohashi, The KEAP1-NRF2 System: a Thiol-Based Sensor-Effector Apparatus for Maintaining Redox Homeostasis. Physiol Rev, 2018. 98(3): p. 1169–1203.

6. Suzuki, T., J. Takahashi, and M. Yamamoto, Molecular Basis of the KEAP1-NRF2 Signaling Pathway. Mol Cells, 2023. 46(3): p. 133–141.

7. Gacesa, R., W.C. Dunlap, D.J. Barlow, R.A. Laskowski, and P.F. Long, Rising levels of atmospheric oxygen and evolution of Nrf2. Sci Rep, 2016. 6: p. 27740.

8. Harris, I.S. and G.M. DeNicola, The Complex Interplay between Antioxidants and ROS in Cancer. Trends Cell Biol, 2020. 30(6): p. 440–451.

9. Cloer, E.W., D. Goldfarb, T.P. Schrank, B.E. Weissman, and M.B. Major, NRF2 Activation in Cancer: From DNA to Protein. Cancer Res, 2019. 79(5): p. 889–898.

10. Pillai, R., M. Hayashi, A.M. Zavitsanou, and T. Papagiannakopoulos, NRF2: KEAPing Tumors Protected. Cancer Discov, 2022. 12(3): p. 625–643.

11. Hamad, S.H., et al., TP53, CDKN2A/P16, and NFE2L2/NRF2 regulate the incidence of pure- and combined-small cell lung cancer in mice. Oncogene, 2022. 41(25): p. 3423–3432.

12. Hamad, S.H., et al., NRF2 Activation in Trp53;p16-deficient Mice Drives Oral Squamous Cell Carcinoma. Cancer Res Commun, 2024. 4(2): p. 487–495.

13. Bowman, B.M., et al., A conditional mouse expressing an activating mutation in NRF2 displays hyperplasia of the upper gastrointestinal tract and decreased white adipose tissue. J Pathol, 2020. 252(2): p. 125–137.

14. DeBlasi, J.M., et al., Distinct Nrf2 Signaling Thresholds Mediate Lung Tumor Initiation and Progression. Cancer Res, 2023. 83(12): p. 1953–1967.

15. Rojo de la Vega, M., E. Chapman, and D.D. Zhang, NRF2 and the Hallmarks of Cancer. Cancer Cell, 2018. 34(1): p. 21–43.

16. Dinkova-Kostova, A.T. and I.M. Copple, Advances and challenges in therapeutic targeting of NRF2. Trends Pharmacol Sci, 2023. 44(3): p. 137–149.

17. Ren, D., et al., Brusatol enhances the efficacy of chemotherapy by inhibiting the Nrf2- mediated defense mechanism. Proc Natl Acad Sci U S A, 2011. 108(4): p. 1433–8.

18. Tsuchida, K., et al., Halofuginone enhances the chemo-sensitivity of cancer cells by suppressing NRF2 accumulation. Free Radic Biol Med, 2017. 103: p. 236–247.

19. Paiboonrungruang, C., et al., Small molecule screen identifies pyrimethamine as an inhibitor of NRF2-driven esophageal hyperplasia. Redox Biol, 2023. 67: p. 102901.

20. Heppler, L.N., et al., The antimicrobial drug pyrimethamine inhibits STAT3 transcriptional activity by targeting the enzyme dihydrofolate reductase. J Biol Chem, 2022. 298(2): p. 101531.

21. Liu, H., et al., Antimalarial Drug Pyrimethamine Plays a Dual Role in Antitumor Proliferation and Metastasis through Targeting DHFR and TP. Mol Cancer Ther, 2019. 18(3): p. 541–555.

22. Brown, J.I., et al., Investigating the anti-cancer potential of pyrimethamine analogues through a modern chemical biology lens. Eur J Med Chem, 2024. 264: p. 115971.

23. Brown, J.R., et al., Targeting constitutively active STAT3 in chronic lymphocytic leukemia: A clinical trial of the STAT3 inhibitor pyrimethamine with pharmacodynamic analyses. Am J Hematol, 2021. 96(4): p. E95–e98.

24. Manoharan, S. and L. Ying Ying, Pyrimethamine reduced tumour growth in pre-clinical cancer models: a systematic review to identify potential pre-clinical studies for subsequent human clinical trials. Biol Methods Protoc, 2024. 9(1): p. bpae021.

25. Cohen, A.A., et al., Dynamic proteomics of individual cancer cells in response to a drug. Science, 2008. 322(5907): p. 1511–6.

26. Wamsley, N.T., et al., Targeted Proteomic Quantitation of NRF2 Signaling and Predictive Biomarkers in HNSCC. Mol Cell Proteomics, 2023. 22(11): p. 100647.

27. Lazear, M.R., Sage: An Open-Source Tool for Fast Proteomics Searching and Quantification at Scale. J Proteome Res, 2023. 22(11): p. 3652–3659.

28. Cox, J. and M. Mann, MaxQuant enables high peptide identification rates, individualized p.p.b.-range mass accuracies and proteome-wide protein quantification. Nature Biotechnology, 2008. 26(12): p. 1367–1372.

29. Cox, J., et al., Accurate proteome-wide label-free quantification by delayed normalization and maximal peptide ratio extraction, termed MaxLFQ. Mol Cell Proteomics, 2014. 13(9): p. 2513–26.

30. Tyanova, S., et al., The Perseus computational platform for comprehensive analysis of (prote)omics data. Nature Methods, 2016. 13(9): p. 731–740.

31. Perez-Riverol, Y., et al., The PRIDE database at 20 years: 2025 update. Nucleic Acids Res, 2024.

32. Adams, K.J., et al., Skyline for Small Molecules: A Unifying Software Package for Quantitative Metabolomics. Journal of Proteome Research, 2020. 19(4): p. 1447–1458.

33. Lazzara, P.R., et al., Synthesis and Evaluation of Noncovalent Naphthalene-Based KEAP1-NRF2 Inhibitors. ACS Med Chem Lett, 2020. 11(4): p. 521–527.

34. Ercikan-Abali, E.A., et al., Dihydrofolate reductase protein inhibits its own translation by binding to dihydrofolate reductase mRNA sequences within the coding region. Biochemistry, 1997. 36(40): p. 12317–22.

35. Ducker, G.S. and J.D. Rabinowitz, One-Carbon Metabolism in Health and Disease. Cell Metab, 2017. 25(1): p. 27–42.

36. Wu, K., A.E. El Zowalaty, V.I. Sayin, and T. Papagiannakopoulos, The pleiotropic functions of reactive oxygen species in cancer. Nat Cancer, 2024. 5(3): p. 384–399.

37. Hayes, J.D., A.T. Dinkova-Kostova, and K.D. Tew, Oxidative Stress in Cancer. Cancer Cell, 2020. 38(2): p. 167–197.

38. Zhou, X., et al., Pyrimethamine Elicits Antitumor Effects on Prostate Cancer by Inhibiting the p38-NF-κB Pathway. Frontiers in Pharmacology, 2020. 11.

39. Ramchandani, S., et al., The multifaceted antineoplastic role of pyrimethamine against human malignancies. IUBMB Life, 2022. 74(3): p. 198–212.

40. Kitamura, H., Y. Onodera, S. Murakami, T. Suzuki, and H. Motohashi, IL-11 contribution to tumorigenesis in an NRF2 addiction cancer model. Oncogene, 2017. 36(45): p. 6315–6324.

41. Okazaki, K., T. Papagiannakopoulos, and H. Motohashi, Metabolic features of cancer cells in NRF2 addiction status. Biophys Rev, 2020. 12(2): p. 435–441.

42. Pouremamali, F., A. Pouremamali, M. Dadashpour, N. Soozangar, and F. Jeddi, An update of Nrf2 activators and inhibitors in cancer prevention/promotion. Cell Commun Signal, 2022. 20(1): p. 100.

43. Ramchandani, S., The multifaceted antineoplastic role of pyrimethamine against human malignancies. IUBMB life, 2021. 74(3): p. 198–212.

44. Saadat, F., et al., Effect of pyrimethamine in experimental rheumatoid arthritis. Med Sci Monit, 2005. 11(8): p. Br293-9.

45. Raimondi, M.V., et al., DHFR Inhibitors: Reading the Past for Discovering Novel Anticancer Agents. Molecules, 2019. 24(6).

46. Brown, J.I., et al., Investigating the anti-cancer potential of pyrimethamine analogues through a modern chemical biology lens. European Journal of Medicinal Chemistry, 2024. 264: p. 115971.

47. Bedrossian, C.W., W.C. Miller, and M.A. Luna, Methotrexate-induced diffuse interstitial pulmonary fibrosis. South Med J, 1979. 72(3): p. 313–8.

48. Silverman, E., et al., Leflunomide or methotrexate for juvenile rheumatoid arthritis. N Engl J Med, 2005. 352(16): p. 1655–66.

49. Albrecht, K. and U. Müller-Ladner, Side effects and management of side effects of methotrexate in rheumatoid arthritis. Clin Exp Rheumatol, 2010. 28(5 Suppl 61): p. S95–101.

50. Zachariae, H., Methotrexate side-effects. Br J Dermatol, 1990. 122 **Suppl 36**: p. 127–33.

51. Maksimovic, V., et al., Molecular mechanism of action and pharmacokinetic properties of methotrexate. Mol Biol Rep, 2020. 47(6): p. 4699–4708.

52. Muller, R.B., J. von Kempis, S.R. Haile, and M.H. Schiff, Effectiveness, tolerability, and safety of subcutaneous methotrexate in early rheumatoid arthritis: A retrospective analysis of real-world data from the St. Gallen cohort. Semin Arthritis Rheum, 2015. 45(1): p. 28–34.

53. Daraghmeh, D.N., C. King, and M.D. Wiese, A review of liquid biopsy as a tool to assess epigenetic, cfDNA and miRNA variability as methotrexate response predictors in patients with rheumatoid arthritis. Pharmacol Res, 2021. 173: p. 105887.

54. Bluett, J., et al., Risk factors for oral methotrexate failure in patients with inflammatory polyarthritis: results from a UK prospective cohort study. Arthritis Res Ther, 2018. 20(1): p. 50.

